# New T2T assembly of *Cryptosporidium parvum* IOWA annotated with reference genome gene identifiers

**DOI:** 10.1101/2023.06.13.544219

**Authors:** Rodrigo de Paula Baptista, Rui Xiao, Yiran Li, Travis C. Glenn, Jessica C. Kissinger

## Abstract

*Cryptosporidium parvum* is a significant pathogen causing gastrointestinal infections in humans and animals, that is spread through the ingestion of contaminated food and water. Despite its global impact on public health, generating a *C. parvum* genome sequence has always been challenging due to a lack of *in vitro* cultivation systems and challenging sub-telomeric gene families. A gapless telomere to telomere genome assembly has been created for *Cryptosporidium parvum* IOWA obtained from Bunch Grass Farms, named here as CpBGF. There are 8 chromosomes that total 9,259,183 bp. The new hybrid assembly which was generated with Illumina and Oxford Nanopore resolves complex sub-telomeric regions of chromosomes 1, 7 and 8. To facilitate ease of use and consistency with the literature, whenever possible, chromosomes have been oriented and genes in this annotation have been given the same gene IDs used in the current reference genome sequence generated in 2004. The annotation of this assembly utilized considerable RNA expression evidence, thus, untranslated regions, long noncoding RNAs and antisense RNAs are annotated. The CpBGF genome assembly serves as a valuable resource for understanding the biology, pathogenesis, and transmission of *C. parvum*, and it facilitates the development of diagnostics, drugs, and vaccines against cryptosporidiosis.

## Background & Summary

*Cryptosporidium parvum* is an apicomplexan parasite that infects the gastrointestinal tract of humans and animals. It is a leading cause of waterborne disease outbreaks worldwide and is responsible for significant morbidity and mortality in immunocompromised individuals and infants ^1^. *Cryptosporidium* infection occurs through the ingestion of contaminated water, food, or fecal matter, making it a significant public health concern ^2^.

Cryptosporidiosis is a neglected disease that has historically been underdiagnosed and underreported ^3^. Although improvements in diagnostic techniques have led to increased detection and reporting of cases, the true burden of disease is likely underestimated due to a lack of surveillance in many countries. The disease is particularly concerning in developing countries, where access to clean water and sanitation is limited, and outbreaks are common. However, in the U.S., cryptosporidiosis is one of the major waterborne diseases related to diarrhea and death in children and immunocompromised patients ^4^.

Despite the importance of *Cryptosporidium* as a human and animal pathogen, sequencing its genome has been challenging due to its repetitive subtelomeric ends, and difficulty to obtain enough material for DNA extraction, since there is a lack of *in vitro* cultivation systems to grow these parasites ^5^. Currently, at NCBI there are > 50 whole genome assemblies available for the genus, representing ∼ 15 species. However, only the *Cryptosporidium parvum* IOWA genome sequence has chromosomal physical mapping information available ^6^. Due to the challenges with material and the extensive use of short-read sequencing, most assemblies are fragmented and incomplete, limiting the community’s ability to accurately compare genome sequences and study their link to the parasite’s biology, virulence mechanisms and transmission dynamics ^7^.

Recent advances in sequencing technologies have enabled the production of high-quality genome assemblies, which can provide an almost complete, if not complete representation of the genome sequence in the absence of physical mapping information ^8,9^. However, some regions of the genome sequence are still challenging including the sub-telomeric regions of chromosomes 1, 7 and 8 which appear to have been replicated among these three chromosomes and also contain important genes such as copies of 18S and 28S rRNA genes, tryptophan synthase Beta and the MEDLE genes ^8^. With Deep sequencing and a combination of recent bioinformatics approaches, we present the first telomere-to-telomere assembly of the *C. parvum* IOWA-BGF (CpBGF) genome, which includes placement of all subtelomeric regions. Importantly, this new genome assembly mirrors the current reference genome assembly ^6^ including chromosomal orientation and gene naming to facilitate comparisons to the reference genome and existing literature. The new assembly has also resolved the unresolved regions in chromosomes 2, 4 and 5 due to lack of physical mapping to support scaffolding found in the original *C. parvum* IOWAII strain (CpIOWA) reference assembly. The new assembly also resolved all eight telomeric regions, and the annotation contains new ncRNA information including lncRNAs and sncRNAs ^10,11^ and UTR boundaries have been added based on RNA long-read data.

The availability of a complete genome assembly for the *C. parvum* IOWAII strain maintained by Bunch Grass Farms (CpBGF) will facilitate studies aimed at understanding the biology, pathogenesis, and transmission of this important pathogen. It will also provide a valuable resource for the development of new diagnostics, drugs, and vaccines against cryptosporidiosis.

## Methods

### Oocyst material

The oocysts for CpBGF were obtained from a commercial source (Bunch Grass Farms, Deary, ID, USA). A total of 10^8^ oocysts were incubated on ice for 10 min in household bleach (diluted 1 : 4 in water) and then washed twice with cold phosphate-buffered saline (PBS). Excystation of sporozoites was then induced by incubating oocysts in 0.8 % sodium taurodeoxycholate (Sigma) in PBS at 37 °C for 1 h. To maintain the potential for high molecular weight (HMW) for long read sequencing, sporozoite genomic DNA was extracted using the traditional phenol-chloroform DNA extraction method ^12^.

### Whole Genome Sequencing and assembly

Illumina reads were retrieved from NCBI’s sequence read archive SRR11516703. These reads were generated on a MiSeq Illumina platform with 300 bp paired-end reads from Bunch Grass Farm oocysts collected in 2017. Illumina sequencing data were quality-checked and trimmed using Trimmomatic v0.39 ^13^.

ONT library preparation was performed using the SQK-LSK109 Ligation Sequencing Kit and the Rapid Barcoding Sequencing Kit (Oxford Nanopore Technologies, Oxford, UK) following the manufacturer’s instructions. The library was sequenced on an ONT MinION device with R9.4.1 flow cells.

The genome assembly was performed using two Long-read assembly approaches: (i) Flye v2.8.2 ^14^; and (ii) NECAT v0.3.3 ^15^. Both assemblies were evaluated, and the structures were compared to identify any regions that differed. NextPolish v2.3.0 ^16^ was used to correct base call errors in the assembly using the CpBGF short read sequencing data. The NECAT genome was chosen as the reference model since it was able to better resolve the telomeric regions, and Geneious prime (v2019) was used to manually curate potential regions that needed to be corrected. Those structural corrections were all supported by long read alignments generated using Minimap2 v2.24 ^17^. Genome statistics were calculated using QUAST v5.0.2 ^18^. Telomeres were detected using the python script FindTelomeres (https://github.com/JanaSperschneider/FindTelomeres) looking for patterns at the start and end of the contigs only. The sequence of the *Cryptosporidium* telomere repeat is “CCTAAA/ AGGTTT”, but the detection also works with the default settings available in the original script.

### Genome annotation

Genome annotation was performed in two ways: (i) by transferring gene annotations based on homology using Liftoff v1.6.3 ^19^ using the *C. parvum* IOWA-ATCC (CpIA) genome as a reference; and (ii) by generating an *ab initio* prediction with BRAKER2 ^20^ trained using a mix of available CpBGF RNA-seq data as was done previously for CpIOWA-ATCC ^8^. Thus, the pipeline integrates multiple evidence-based approaches, including *ab initio* gene prediction based on transcriptomic models, and homology-based prediction to annotate genes. Both annotation tracks were added to a WebApollo2 environment ^21^ for manual curation.

### Gene structure and UTR length curation using Iso-seq

Two major issues for *Cryptosporidium* genome annotation are the detection of untranslated region (UTR) boundaries and validation of annotated gene structures. It is difficult because *C. parvum*’s highly compact genome has an average of only 500 bp of intergenic space between the stop codon of one gene and the start codon of the next. This compactness causes frequent UTR overlap, which makes it difficult to differentiate gene boundaries using short-read RNA-seq techniques. To solve this problem and generate more reliable UTR annotation predictions for this new annotation, we sequenced sporozoite total RNA using PacBio Iso-Seq and ONT direct RNAseq methods. Iso-Seq data were processed using SMRT LINK v11.0 (PacBio, CA) and TAMA ^22^. First, raw reads were quality filtered to remove sequences with either incomplete read cycle or lacking a poly-A tail. Filtered circular consensus sequencing (CCS) reads are subsequently trimmed of adapters and poly-A tails, then collapsed into linear full-length long reads with PacBio’s SMRTlink tools. Reads are mapped to the CpBGF genome using minimap2 v.2.24 ^17^. The mapped reads are then fed into the TAMA pipeline to be collapsed into transcript models using TAMA’s high stringency mode. ORFs were predicted and assigned to individual transcript models using NCBI BLAST. The TAMA pipeline generated transcript models were used in conjunction with transcript models obtained from the annotation pipeline to update the gene model structures including UTRs. The UTR boundaries were determined collectively using both Iso-seq and short-read stranded RNA-seq with a boundary cutoff requiring > 90% read support.

These approaches helped generate full-length transcripts of all expressed genes, facilitating the genome annotation and determination of UTR boundaries. These tracks were also added to a WebApollo 2 instance for use in manual curation.

### Gene function annotation

Gene function annotation was performed by comparing the predicted protein sequences to previous predictions made for CpIA ^8^, which was supported by protein sequence similarity using BLAST against Swiss-Prot, Trembl, and the NCBI nonredundant protein database, I-TASSER v5.1 ^23^ and Interproscan v5 ^24^. The best hit with an E-value cutoff of 1E-5 or better was used to assign putative functions to the genes. This approach benefits from the previous annotation and updates generated by the *Cryptosporidium* community and submitted to repositories.

Non-coding RNA annotation was performed using tRNA-scan-SE v2.0.7 ^25^ and Infernal v1.1.3 ^26^ which can detect most of the known ncRNA families such as tRNA’s and rRNAs. Additionally, *Cryptosporidium* long non-coding and small non-coding RNAs described by Li et al. ^10,11^ were added.

### Comparison with previous *C. parvum* assemblies

Structural comparative analysis was performed using Minimap2 v2.24 ^17^, Samtools v.1.16.1^27^ and Progressive MAUVE v2.4.0 ^28^ to identify syntenic regions among the available chromosomal level assemblies. Gene orthology was determined using OrthoFinder v2.5.4 ^29^ to identify orthologous genes and detect potential duplication events. The orthology information was also useful to generate matching gene IDs to the original CpIOWAII genome annotation that has been used as a reference by the community for the past 18 years. We believe that using matching ID numbers will benefit the community.

To analyze single nucleotide variants (SNVs) across the available strains, we used the Genome Analysis Toolkit v.4.1.4.0 (GATK) ^30^. This software was chosen for its ability to detect small variants such as SNPs and indels with high sensitivity and specificity. We first aligned the reads from different available *C. parvum* IOWA strains to the new CpBGF genome sequence using the BWA-MEM v0.7.12 ^31^ alignment tool and then GATK to call variants across all samples. We then compared the called variants across the different strains to identify and compare the differences and similarities. All called variants were than submitted to a variant annotation pipeline using snpEFF v5.1 ^32^. The results from this analysis are used to assess the suitability of the new genome sequence for the community with regards to variability across strains.

## Results

The hybrid CpBGF genome assembly resulted in a genome sequence consisting of 8 contigs with an N50 value of 1,107,426 bp, and a genome length of 9,259,183 bp (Table 1). The genome sequence has a GC content of 30.04%, and the BUSCO evaluation showed that the genome assembly captured 96.2% of the expected apicomplexan genes, suggesting that the genome sequence is likely complete. *C. parvum* has lost many genes still present in other apicomplexans. The analysis revealed that the CpBGF is the first telomere-to-telomere genome assembly for *C. parvum*.

**Table 1.** Genome Statistics of the available chromosomal level *C. parvum* genome assemblies.

| Statistics | CpIOWAll<sup>a</sup> | CpBGF | CpIOWA-ATCC<sup>b</sup> | CpBCM-BGF<sup>c</sup> |
| --- | --- | --- | --- | --- |
| **# of contigs** | 8 | 8 | 8 | 8 |
| **Largest contig** | 1,344,712 | 1,379,419 | 1,332,634 | 1,355,650 |
| **Total length** | 9,102,324 | 9,259,183 | 9,122,263 | 9,197,619 |
| **N50** | 1,104,417 | 1,107,426 | 1,108,396 | 1,108,772 |
| **L50** | 4 | 4 | 4 | 4 |
| **GC%** | 30.22 | 30.04 | 30.18 | 30.11 |
| **N's** | 18,558 | 0 | 0 | 0 |
| **# of assembled telomeres** | 3 | 16 | 13 | 12 |
| **Annotation contributing to transcripts (bp)** | 7,430,056 | 8,203,418* | 7,340,634 | NA |

<sup>a</sup> (Abrahamsen et al., 2004)<sup>6</sup>; <sup>b</sup> (Baptista et al., 2022)<sup>8</sup>; <sup>c</sup> (Menon et al., 2022)<sup>9</sup> \*ncRNA information and UTRs are added compared to previous versions; NA, Not Available, Genome sequence is unannotated.

Comparative genomic analysis with other *C. parvum* genome sequences revealed high conservation of the genome structure among the new long-read-driven *C. parvum* assemblies. Other than the genome size, the major structural difference between CpBGF and the other publicly available long-read generated genome assemblies is the completeness of telomere assembly, e.g., CpIOWA-ATCC ^8^ and CpBCM-BGF ^9^ (Table 1).

When comparing all four *C. parvum* genome sequences for synteny, besides their high similarity, we can observe that the CpBGF genome sequence follows the same chromosome level orientation as the original genome assembly (Figure 1). However, the inversions that exist in the original CpIOWAII genome assembly on chromosomes 2, 4 and 5 are not observed in any of the long-read assemblies. The CpBGF genome assembly thus addresses orientation discrepancies relative to the original assembly and also keeps a similar gene ID based on homology to genes in the original assembly, which facilitates linkage to published gene IDs. For example, the commonly used *gp60* gene, reference gene ID cgd6_1080 will be either cpbgf6_1080 or cpbgf_6001080 depending on the outcome of negotiations with GenBank (they no longer allow chromosome numbers in the prefix).

**Figure 1.**
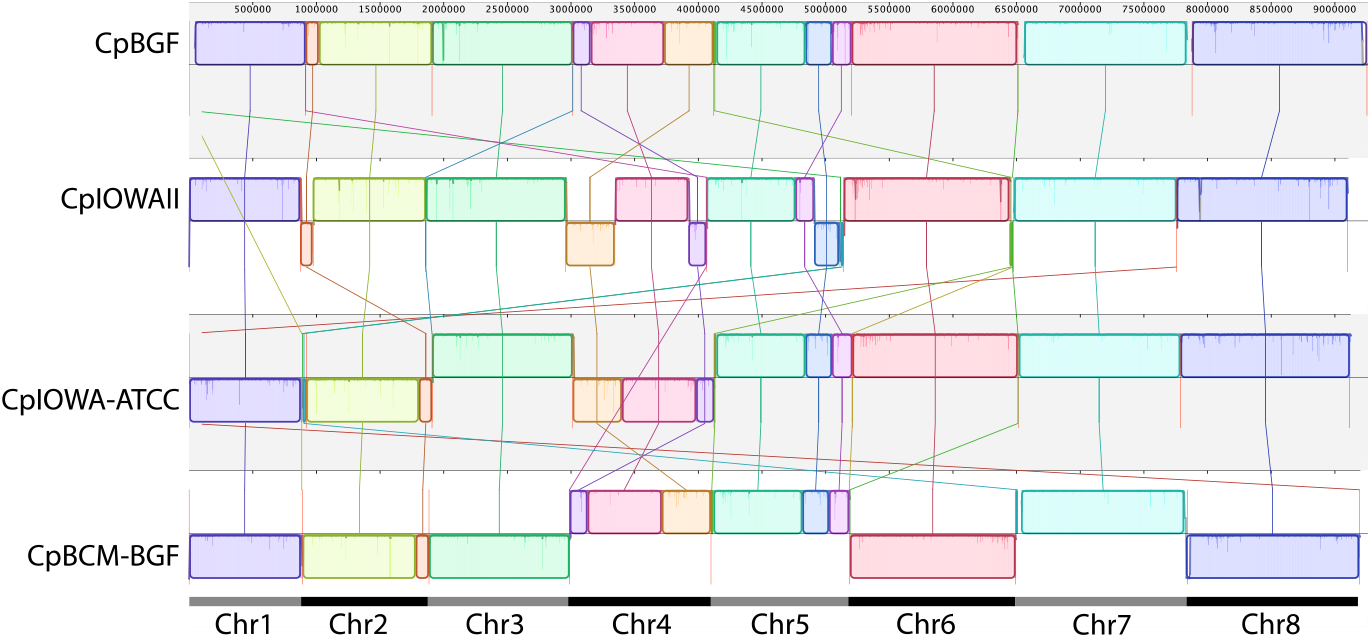
Synteny plot between the new CpBGF genome sequence and other *C. parvum* chromosomal level genome sequence. Progressive Mauve synteny plot showing discrepancies in the chromosomal order of CpIOWA-ATCC and CpBCM-BGF when compared to CpIOWAII. Chr2, Chr4 and Chr5 in CpIOWAII have inverted and translocated regions relative to the assemblies generated for other *C. parvum* IOWA isolates. Note Chrs 2, 4 and 5 have been colored in the same colors as the discontiguous scaffolded contigs in CpIOWAII but they are gapless, contiguous sequences in the long-read assemblies.

The sub-telomeric regions of Chr1, 7, and 8, which were challenging to sequence in previous studies ^8^, are well-represented in the new assembly. The results demonstrate the usefulness of the BGF strain as a reliable reference genome for *C. parvum*.

All genome sequences present a *gp60* type IIa, but an analysis of their amino acidic sequences reveals see small differences between them. Most differences are related to the CpIOWAII assembly and may be caused by accumulated differences over time in the current extant isolates or sequencing limitations at the time of the original assembly. CpBCM-BGF also shows an amino acid difference in Lysine (K) instead of a Threonine (T) at position 296 (Figure 2).

**Figure 2.**
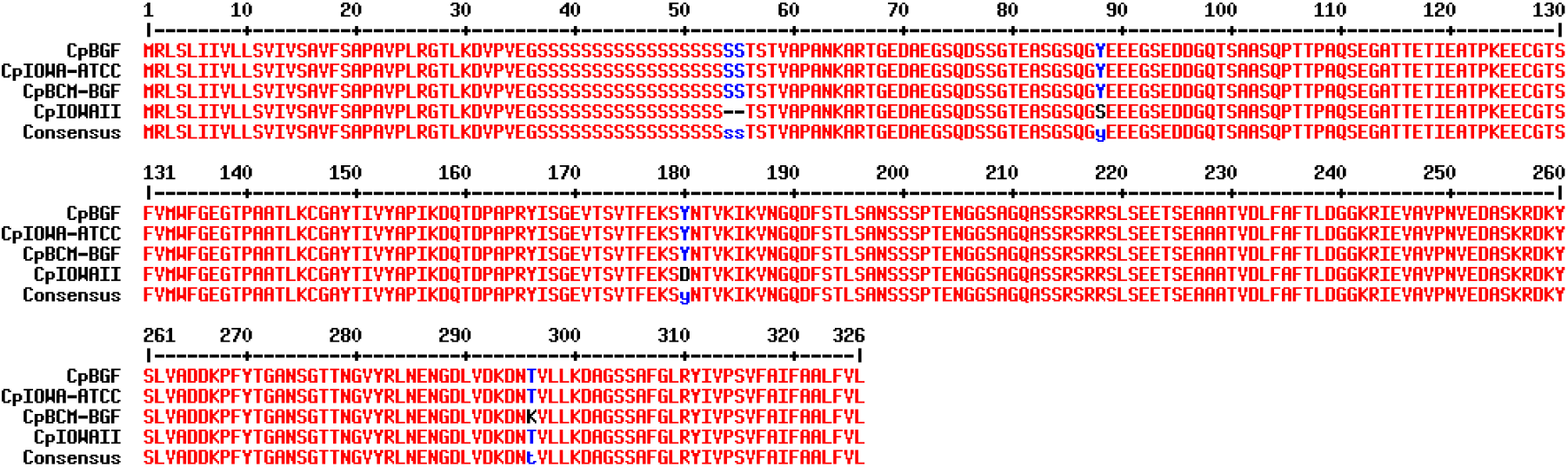
Amino acid alignment between GP60 sequences from all four *C. parvum* genome sequences analyzed.

To assess variation in gene content between the genome assemblies an orthology analysis was performed. Orthology analysis was performed using OrthoFinder to identify genes shared by the four isolates. A total of 15,597 annotated genes used in this analysis are shared by at least two assemblies. CpIOWAII has the largest number of singletons mostly related to sequences not matching the updated annotated genes and missing regions consisting of gaps. Clustering analysis of the annotated genes revealed a close relationship between all strains, as expected (Figure 3). CpBGF and CpBCM-BGF are the most similar, when compared to IOWA-ATCC (CpIA), and most differences are due to updates in the evidence-based genome annotation of CpBGF. Since CpBCM-BGF didn’t have an available annotation, we transferred the annotation based on CpBGF using liftoff to make the comparisons. The only differences between their gene content is related to the missing subtelomeric regions in CpBCM-BGF. Overall, the orthology analysis provided insights into both the assembly completeness and the genetic relatedness of the *C. parvum* IOWA samples. There are differences between the original CpIOWA reference and the newly generated genome sequences.

**Figure 3.**
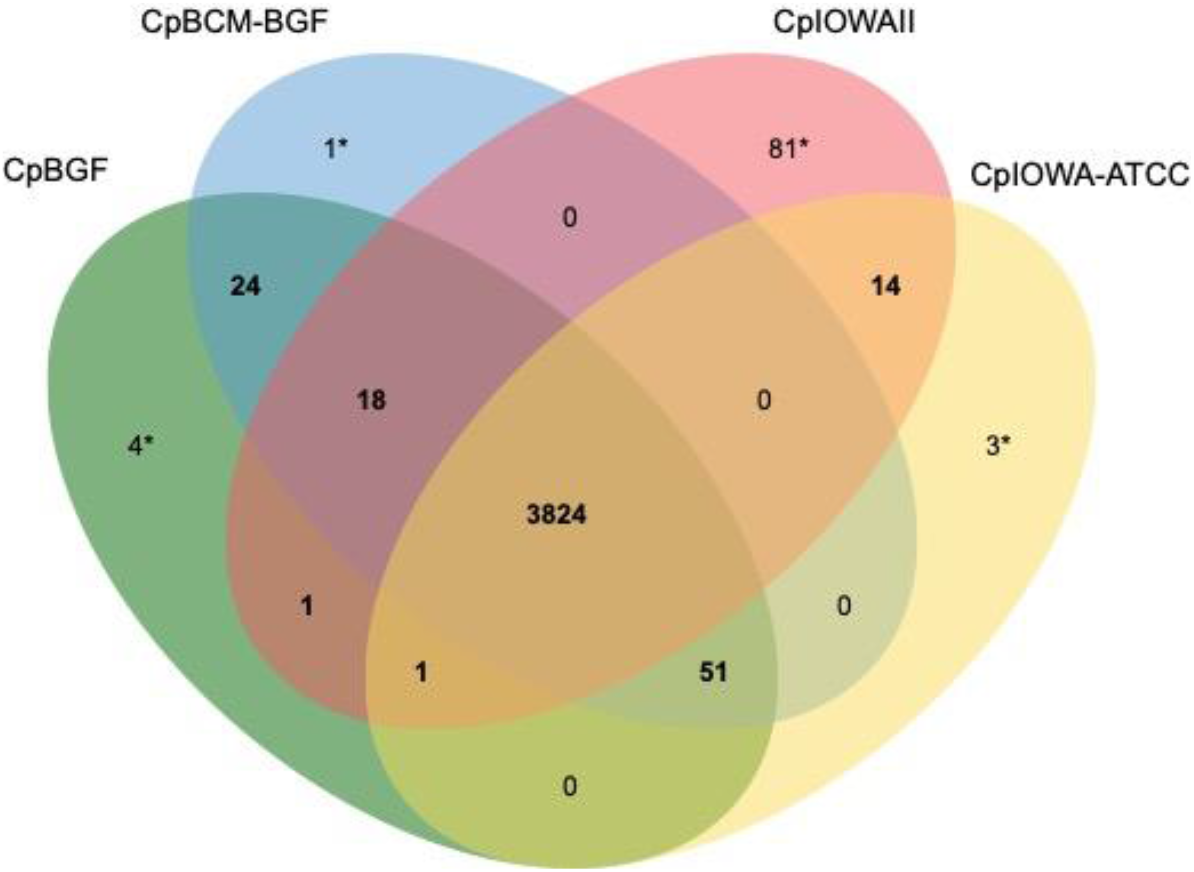
Venn diagram indicating the number of shared Orthologue groups. Singletons observed between CpBCM-BGF and CpBGF are related to missing subtelomeric regions in CpBCM-BGF. *Singletons – Protein with no shared ortholog found due to assembly limitations and variation.

Variant analysis revealed a high degree of conservation among the four genome sequences with only a few variants present when compared to CpBGF Illumina short-reads. The distribution of SNPs across the genome showed a low number of SNPs, with most regions exhibiting no variation (Figure 4A). The variants found in the genome were all found in intergenic regions in CpBGF and mostly found in intergenic regions and gap driven regions, which are variants located in gaped regions in the CpIOWAII assembly. A total of 16 variants found across 12 genes in CpIOWAII were found to be non-synonymous (Figure 4B; Table S1).

**Figure 4.**
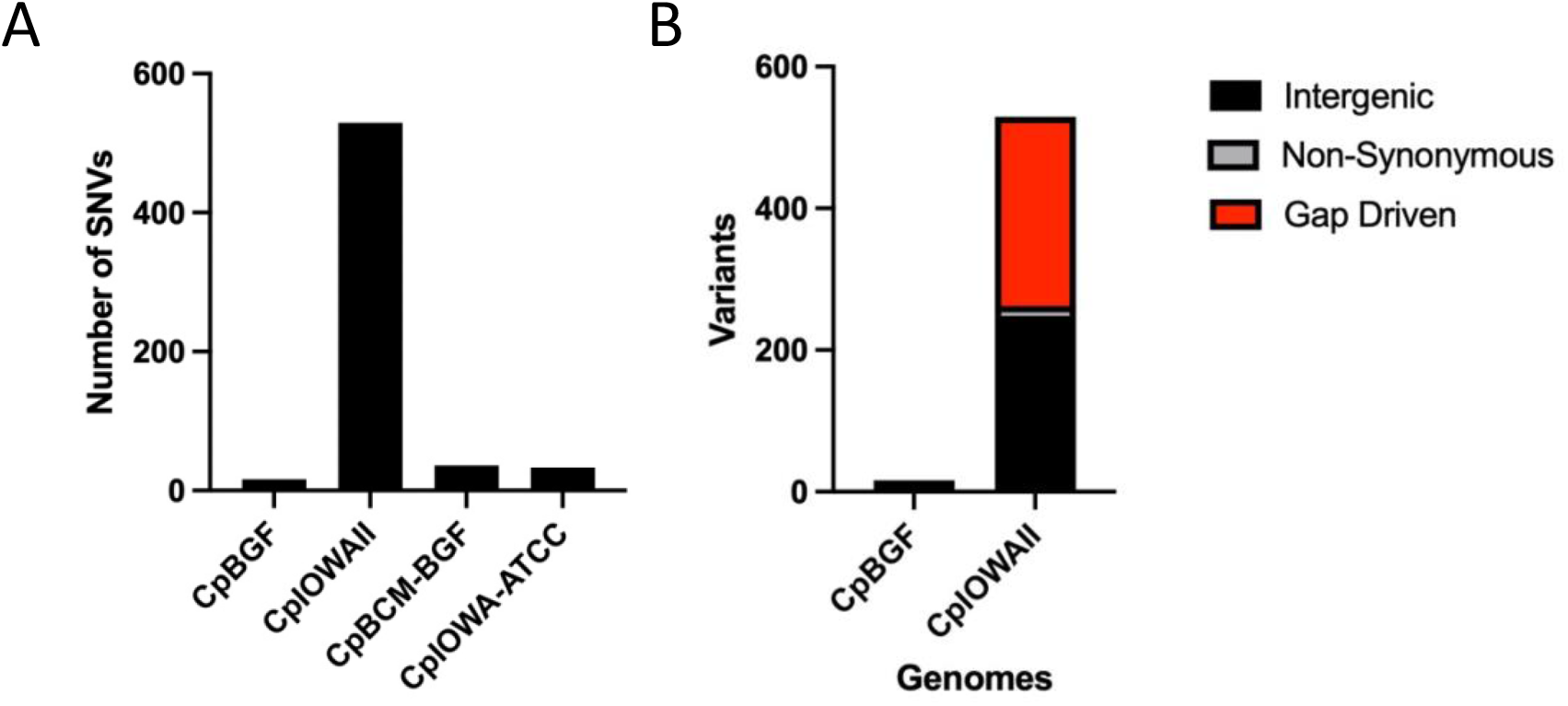
Comparative SNV distribution across all four genomes. (A) total number of variants found when compared to BGF. (B) Variant types found in CpBGF and CpIOWAII separated by genome feature location.

## Discussion

The availability of high-quality reference genomes is essential for studying the biology and pathogenicity of parasites like *Cryptosporidium*. However, obtaining complete genome assemblies of this species is challenging due to their complexities and lack of material for sequencing. Our study addressed these challenges by using a long- and short-read hybrid sequencing approach to generate the first telomere-to-telomere genome assembly for the *C. parvum* IOWA strain obtained from Bunch Grass Farms, BGF, a commercial supplier widely used by the community in the United States. The assembly fills gaps present in the current *C. parvum* IOWAII strain reference assembly and provides full representation of sub-telomeric regions on chromosomes 1, 7, and 8. The differences between the genome assemblies generated thus far reflect technical challenges and advances in sequencing and assembly approaches. There are also true biological differences as this strain, IOWA, which must be maintained by animal passage has already been observed to be evolving ^33^.

The *C. parvum* IOWA-BGF strain genome assembly will be valuable for researchers studying *Cryptosporidium*. The gene IDs provided in this new assembly and annotation follow the former CpIOWAII naming whenever possible thus facilitating ease of use. The availability of a telomere-to-telomere genome assembly has and will continue to enable more comprehensive analyses of the gene repertoire of *C. parvum* and future functional characterization. It will also aid in the identification of virulence factors and other genes involved in pathogenicity and drug resistance

### Data Records

The genomic data for the *Cryptosporidium parvum* IOWA-BGF strain has been deposited in the NCBI database under the BioProject PRJNA983265 and SRA accession SRR13777123. The assembly statistics, BUSCO assessment, Illumina reads coverage, single nucleotide polymorphisms have been made available for public access and download. All relevant information regarding the sequencing methods and analysis pipeline are provided to ensure the reproducibility of the results.

### Technical Validation

To validate the quality of the *Cryptosporidium parvum* IOWA-BGF genome assembly, we used Benchmarking Universal Single-Copy Orthology (BUSCO) software v5.4.6 ^34^ to search Apicomplexa databases containing a total of 446 orthologous single-copy genes. The results showed that 96.0% of the complete single-copy genes were retrieved, of which none were duplicated. Only 0.9% of BUSCO genes were fragmented, and 3.1% were missing from the genome. These results indicate that the genome assembly has high completeness.

To further validate the accuracy of the genome assembly, we aligned the filtered short Illumina reads back to the genome assembly using BBmap v.38.93 software ^35^. Approximately 95% of the short reads mapped to the genome, and 5% were marked as not mapped. The ratios of multiallelic single nucleotide polymorphisms (SNPs) were 0%, indicating that the assembly had high single, base-level accuracy.

Overall, the high BUSCO completeness scores and accurate mapping of Illumina reads to the long-read assembly support the quality and reliability of the new CpBGF genome assembly.

## Code availability

1. Chimeric read detection:
  - Minimap2/yacrd: minimap2 -x ava-ont -g 500 CpBGF_ONT_raw.fastq CpBGF_ONT_raw.fastq > overlap.paf yacrd -i overlap.paf -o reads.yacrd
2. Genome Assembly:
  - NECAT/0.0.1: necat.pl correct necat_config.txt necat.pl assemble necat_config.txt necat.pl bridge necat_config.txt
  - necat_config.txt: PROJECT=BGF_NECAT ONT_READ_LIST=BGF_ONT.txt GENOME_SIZE=9200000 THREADS=5 MIN_READ_LENGTH=3000 PREP_OUTPUT_COVERAGE=30 OVLP_FAST_OPTIONS=-n 500 -z 20 -b 2000 -e 0.5 -j 0 -u 1 -a 1000 OVLP_SENSITIVE_OPTIONS=-n 500 -z 10 -e 0.5 -j 0 -u 1 -a 1000 CNS_FAST_OPTIONS=-a 2000 -x 4 -y 12 -l 1000 -e 0.5 -p 0.8 -u 0 CNS_SENSITIVE_OPTIONS=-a 2000 -x 4 -y 12 -l 1000 -e 0.5 -p 0.8 -u 0 TRIM_OVLP_OPTIONS=-n 100 -z 10 -b 2000 -e 0.5 -j 1 -u 1 -a 400 ASM_OVLP_OPTIONS=-n 100 -z 10 -b 2000 -e 0.5 -j 1 -u 0 -a 400 NUM_ITER=2 CNS_OUTPUT_COVERAGE=30 CLEANUP=1 USE_GRID=false GRID_NODE=0 SMALL_MEMORY=0 POLISH_CONTIGS=true
  - Genome Polishing:
    - NextPolish v.1.2.4 : sgs_options = -max_depth 200, lgs_options = -min_read_len 1k - max_read_len 100k, and lgs_minimap2_options = -x map-ont.
  - Genome Annotation:
    - BRAKER2: braker.pl --species=yourSpecies --genome=genome.fasta \ --rnaseq_sets_ids=SRA_ID1,SRA_ID2 \ --rnaseq_sets_dirs=/path/to/local/fastq/files/ \ --UTR=on --addUTR=on --ab_initio
    - Liftoff: liftoff -g CpIA_reference.gff -o CpBGF_liftoff.gff -copies -cds CpBGF_genome.fasta CpIA_genome.fasta
  - Isoseq analysis:
    - SMRTlink: pbmm2 index cp_bgf.fa cp_bgf.mmi pbmm2 align \ cp_bgf.mmi cp.flnc.bam cp_bgf_isoseq_aligned.bam \ --sort -j 8 -J 8 --preset ISOSEQ \ -G 2500 --log-level INFO
    - TAMA: tama_collapse.py -s tama/ cp_bgf_isoseq_aligned.sam \ -f cp_bgf.fa \ -p tama_cp_bgf \ -x no_cap \ -d merge_dup \ -a 100 -z 100 \ -sj sj_priority -lde 1 -sjt 20
  - BUSCO run: busco -i <genome.fasta> -l ./apicomplexa_odb10 -m genome -o <output>
  - Variant Call:
    - BWA/SAMTOOLS: bwa mem -v 2 -M -t 10 $GENOME $R1 $R2 > $SAMPLE.sam samtools view -b -S -o $SAMPLE.bam $SAMPLE.sam
    - PICARD: java -Xmx2g -classpath “$PICARD” -jar $SortSam INPUT=$SAMPLE.bam OUTPUT=$SAMPLE.s.bam VALIDATION_STRINGENCY=LENIENT SORT_ORDER=coordinate java -Xmx2g -classpath “$PICARD” -jar $MarkDuplicates INPUT=$SAMPLE.s.bam OUTPUT=$SAMPLE.sd.bam METRICS_FILE=$SAMPLE.dedup.metrics REMOVE_DUPLICATES=false VALIDATION_STRINGENCY=LENIENT ASSUME_SORTED=true java -Xmx2g -classpath “$PICARD” -jar $ARreadgroups INPUT=$SAMPLE.sd.bam OUTPUT=$SAMPLE.sdr.bam SORT_ORDER=coordinate RGID=$SAMPLE RGLB=$SAMPLE RGPL=illumina RGPU=$SAMPLE RGSM=$SAMPLE VALIDATION_STRINGENCY=LENIENT java -Xmx2g -classpath “$PICARD” -jar $BuildBamIndex INPUT=$SAMPLE.sdr.bam VALIDATION_STRINGENCY=LENIENT
    - GATK: gatk HaplotypeCaller -R $GENOME -I $SAMPLE.sdrsm.bam -O $SAMPLE.GATK.vcf -ploidy $PLOIDY -stand-call-conf 30 gatk VariantFiltration -R $GENOME -V $SAMPLE.GATK.vcf -O $SAMPLE.GATK_Filtered.vcf -cluster 3 -window 10 -filter “QUAL < 30.0 || DP < 10 || QD < 1.5 || MQ < 25.0” --filter-name “StdFilter” -filter “MQ0 >= 4 && ((MQ0 / (1.0 * DP)) > 0.1)” --filter-name “HARD_TO_VALIDATE” -filter “MQ < 40.0 || FS > 60.0” --filter-name “gatkFilter”

Other commands and pipelines used in data processing were executed using their corresponding default parameters.

## Acknowledgements

This work was funded by NIH R01AI14866 to JCK and TCG.

